# Unsupervised deep learning enables blur-free super-resolution in two-photon microscopy

**DOI:** 10.1101/2024.04.30.591870

**Authors:** Haruhiko Morita, Shuto Hayashi, Takahiro Tsuji, Daisuke Kato, Hiroaki Wake, Teppei Shimamura

## Abstract

We developed an unsupervised deep learning method to simultaneously perform deblurring, super-resolution, and segmentation of two-photon microscopy images. Two-photon microscopy is an excellent technique for non-invasively observing deep biological tissues, but blurring during deep imaging has been a challenge. Conventional deblurring methods have limited performance and are not suitable for deblurring two-photon microscopy images. Moreover, methods that simultaneously perform segmentation, which is usually required in downstream analysis, have not been developed. Therefore, in this method (TENET), we precisely modeled the blur of two-photon microscopy and simultaneously achieved deblurring, super-resolution, and segmentation through unsupervised deep learning. In simulation and experimental data, we achieved deblurring, resolution improvement, and segmentation accuracy surpassing conventional methods. Furthermore, we applied the method to live imaging of microglia, enabling quantitative 3D morphological analysis that was previously difficult. This method allows non-invasive visualization of detailed structures in deep biological tissues, and is expected to lead to a more high-definition understanding of biological phenomena. Future applications to time-series morphological analysis of microglia are anticipated.

## Introduction

3D visualization reveals structural information that cannot be obtained in 2D. Particularly in neuroscience, 3D visualization of neurons and glia has been performed using confocal microscopy ^i^ and light-sheet microscopy ^ii^, elucidating their morphology and dynamics^iiiiv^. Microglia change their morphology depending on the presence or absence of stimuli^v^, and non-invasive imaging of deep tissue has been performed using two-photon microscopy^viviiviiiix^, However, two-photon microscopy suffers from axial blurring, making 3D visualization and downstream analysis difficult^x^.

For live imaging with two-photon microscopy, the images suffer from severe axial blurring, high noise levels, and significant motion artifacts. Conventional deconvolution methods^xixiixiii^ as well as deep learning-based deconvolution approaches have been insufficient in removing the blurring from these challenging images. Deconvolution methods require setting an accurate point spread function (PSF), though measuring the PSF inside living tissues^xivxv^ is difficult and lacks accuracy if not done carefully. To address this issue, supervised deep learning models and unsupervised deep learning models have been proposed. However, when applying these conventional methods to two-photon microscopy images, the following problems arise: Supervised methods require labeled data, but there are few datasets available for two-photon microscopy. To address this problem, supervised deep learning models and unsupervised deep learning models have been proposed. However, when applying these conventional methods to two-photon microscopy images, the following problems arise. For supervised methods^xvixviixviii^, labeled data is required, but there are few datasets available for two-photon microscopy. For unsupervised learning methods ^xixxxxxi^, they assume the isotropy of the structure, which means the axial shape is similar to the horizontal shape. However, for live two-photon microscopy imaging where the angle cannot be changed, this isotropic assumption may not hold, making accurate inference difficult.

Two-photon microscopy live imaging not only suffers from large axial blur but also faces issues with noise level and stack misalignment. Furthermore, when performing downstream analysis using the processed images, segmentation is often required^xxiixxiii^. However, there were no methods that simultaneously perform deblurring, denoising, and segmentation. It was necessary to carry out each process one by one. For these reasons, it was difficult to know the three-dimensional morphology and dynamics of deep microglia in an unstimulated state in vivo. Although the relationship between microglial morphology and disease has been pointed out in two-dimensional analysis and three-dimensional analysis using confocal microscopy^xxivxxvxxvixxvii^, there are few examples of analysis that captures three-dimensional time-series changes in the same individual using two-photon microscopy _xxviii_.

In this study, we constructed a model consisting of a combination of deep learning and generative models and developed a method to simultaneously perform unsupervised deblurring, denoising, and segmentation of two-photon microscopy images. The deep learning model takes a 3D two-photon microscopy image as input and outputs an image that has undergone deblurring, denoising, and segmentation processing. The training of this deblurring model consists of two stages: pre-training and fine-tuning. In the first stage, paired data is created by generating simulation images with randomly placed objects and blurring them using a generative model, and the deep learning model is pre-trained. In the second stage, to improve the deblurring accuracy on real images, the deep learning model and the generative model are fine-tuned by performing reconstruction learning using actual images. As a validation of the model, the results of applying the method to simulation images and microglia images are shown.

We provided a method to perform unsupervised, high-accuracy segmentation and 3D visualization of in vivo microglia without using the assumption of structural isotropy.

The proposed model was able to accurately infer the shape of objects in images captured by two-photon microscopy deep tissue imaging. When the model was applied to microglia data, the results showed that the model can remove blur from the image, improve resolution, and obtain clearer images of cells. This method enables the observation and quantification of the dynamics of 3D cell shapes in in vivo two-photon microscopy imaging, which was previously difficult.

## Results

### Schema of the method

In this study, we propose an autoencoder model to achieve unsupervised deblurring, super-resolution, and segmentation of two-photon microscopy images. Figure 1 shows the learning scheme of the framework.

**Figure 1:**
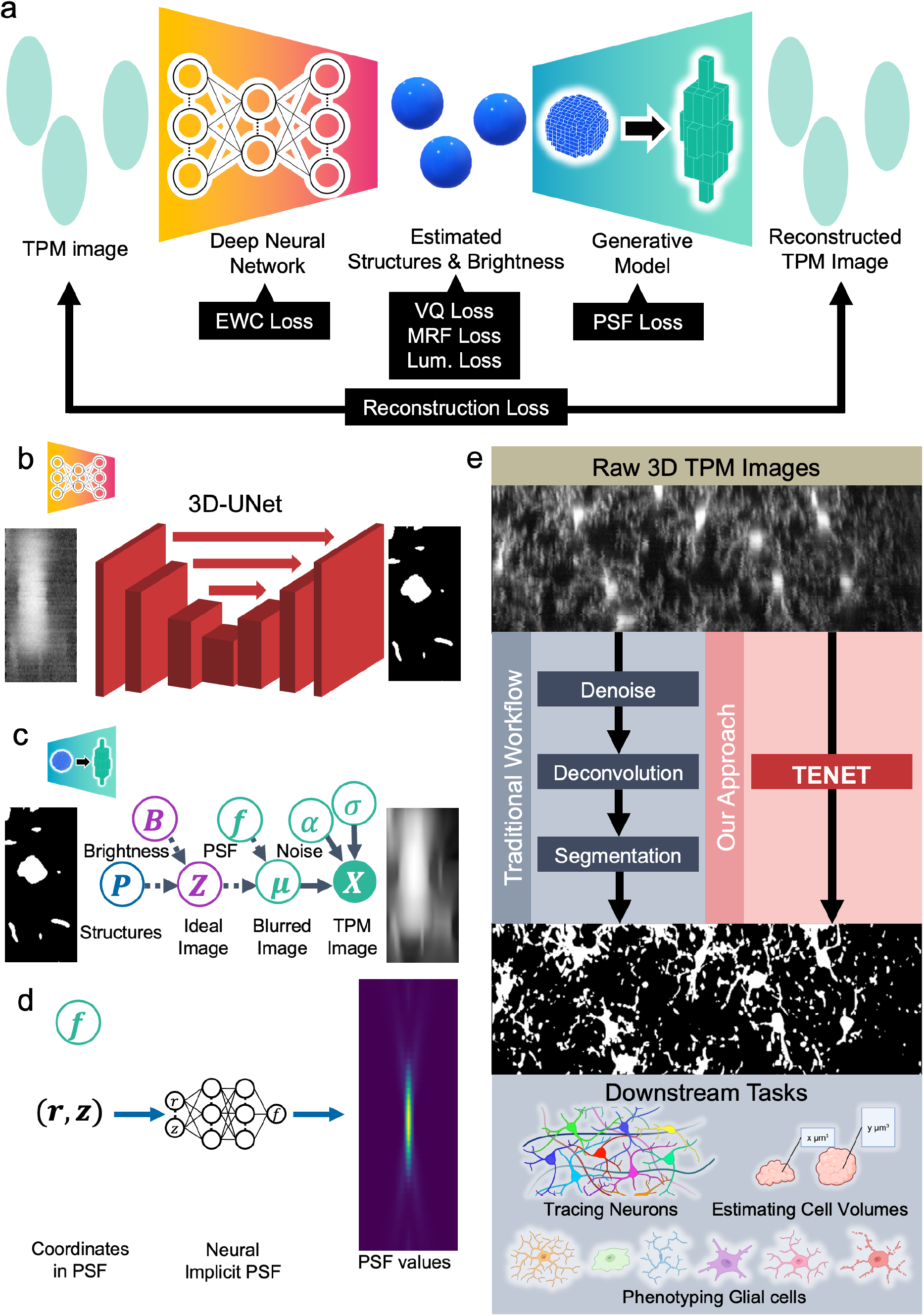
Overview of our framework. **a** Overview of the proposed method. **b** Deblurring model is a deep neural network based on 3D-UNet. **c** Blur generation model simulates the process of image acquisition. **d** Neural implicit PSF is a component of the blur generation model, which represents PSF. **e** Model applications and use cases. Created with BioRender.com.

The framework consists of an encoder from the deep learning model and a decoder from the generative model. The deep learning model infers the deblurred, super-resolved, and segmented image, as well as the brightness, from 3D images such as two-photon microscopy images. The generative model simulates the blurring generation process based on a physically rigorous model.

Our key innovation is to train these two models without actual labeled images by appropriately designing the learning process and loss function.

The learning process in this method is as follows. First, we generate objects with randomly varying brightness and create simulation images blurred by the blur generation model. We use these paired images to pre-train the deblurring model (Figure 1b, c). Next, we pass the actual images through the deblurring model and the blur generation model in sequence and compare the reconstructed images with the original images to fine-tune the two models.

### Experiments with simulation images

For quantitative evaluation, we first applied the model to simulation images (Figure 2a, d). The axial resolution of the images was set to 1/3 of the lateral resolution, and the Gibson-Lanni model^xxixxxx^ was used for the PSF. Experiments were conducted on images with and without Poisson noise added.

**Figure 2:**
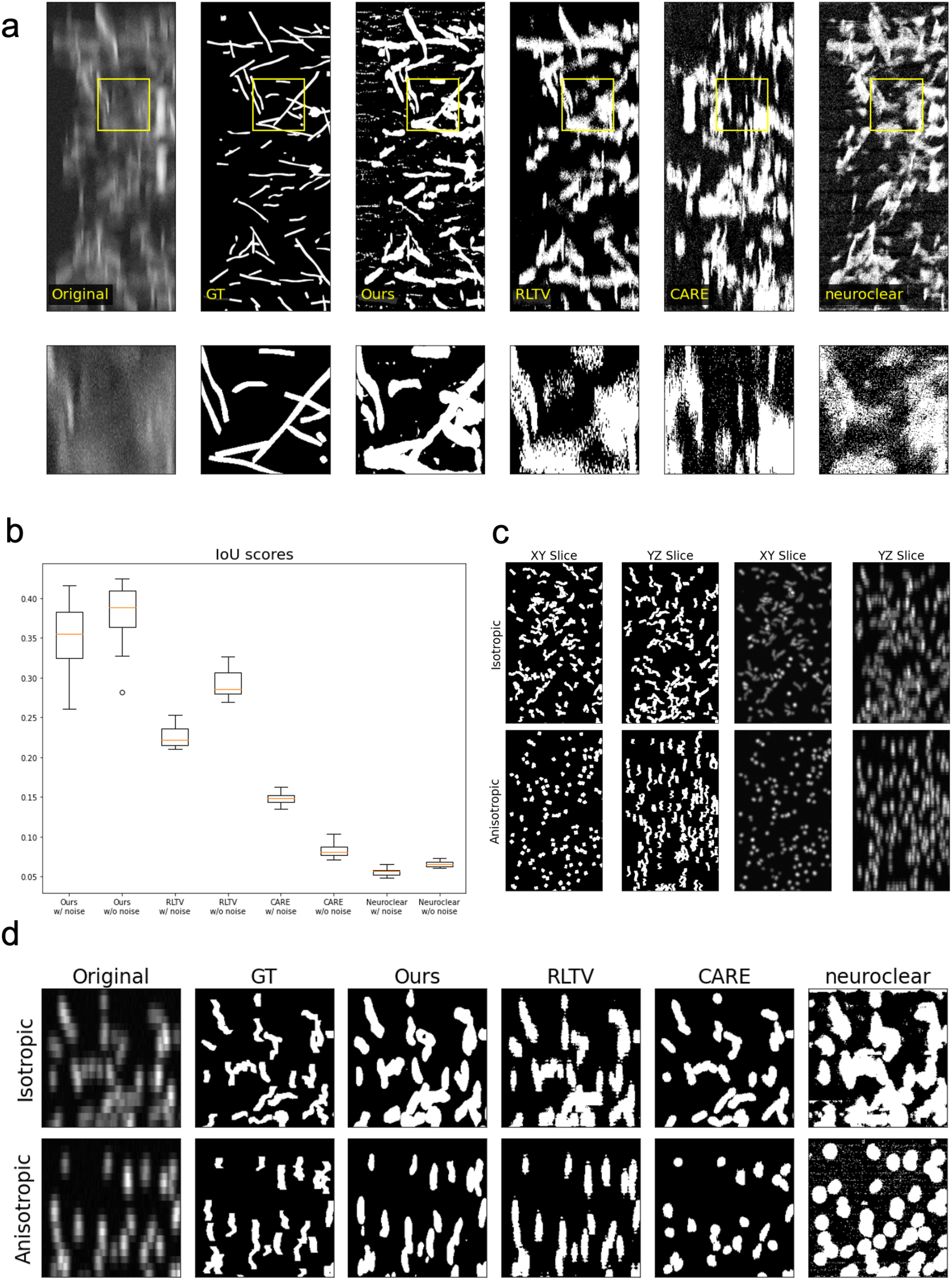
Results with simulation images. **a** Experimental results with simulation images. **b** Comparison of IoU scores with simulation images between proposed method and previous methods. **c** Schema of isotropic/anisotropic simulation volumes. **d** Experimental results with isotropic/anisotropic simulation images, with small Poisson noise.

The model was able to infer the shape of the objects (Figure 2a). Figure 2d shows the IoU (Intersection over Union) between the inference results and the ground truth images. Comparing the results of previous studies and our model’s predictions, our model reconstructs the shape of the objects more accurately than other methods. The deconvolution method RLTV shows reduced accuracy depending on the presence or absence of noise, but our model has a small difference in accuracy and can reconstruct images independently of noise. Regarding CARE and Neuroclear, it is possible that they were unable to properly reconstruct the images because the blur was larger than the size of the input images used in the experiments. In addition to this issue, Neuroclear showed artifacts in regions where nothing was captured in the image, which is thought to be the reason for the low IoU score.

In the next experiment, we verified whether the model could predict structures with anisotropy. Previous studies on unsupervised learning, such as CARE and Neuroclear, assume structural isotropy, meaning that the axial structure is similar to the lateral structure. However, this assumption does not necessarily hold true for two-photon microscopy live imaging, where imaging can only be per-formed from one direction. Our method does not use this assumption, so it is expected to learn even from images of anisotropic structures. To verify this, we randomly generated linear structures on a plane, applied elastic deformation to them, and created simulation images by rotating them randomly or 90 degrees from the center of the stack image along the z-axis (Figure 2c). The randomly rotated simulation images have an isotropic structure, while the 90-degree rotated images have an anisotropic structure. The axial resolution of the images was set to 1/10 of the lateral resolution, and the 3D Gaussian with *σ*_*z*_ = 15, *σ*_*x*_ = 5 (voxels) was used for the PSF. Experiments were conducted on images with small Poisson noise added. Our model and RLTV were able to predict the shape of the objects regardless of isotropy or anisotropy, but CARE and Neuroclear, which rely on structural isotropy for learning, were unable to predict the linear shape in the 90-degree rotated images, and it was found that round artifacts were generated.

### Experiment with actual TPM images

To demonstrate that the method can be applied to biological imaging, we show the results of applying the method to live imaging of microglia in the mouse brain. The original image resolution was 0.31*0.31*1 μm (xy*z), and the field of view was 1024*1024*111 voxels (xy*z). The results of applying the model to this image for deblurring, super-resolution, and segmentation are shown in Figure 3.

**Figure 3:**
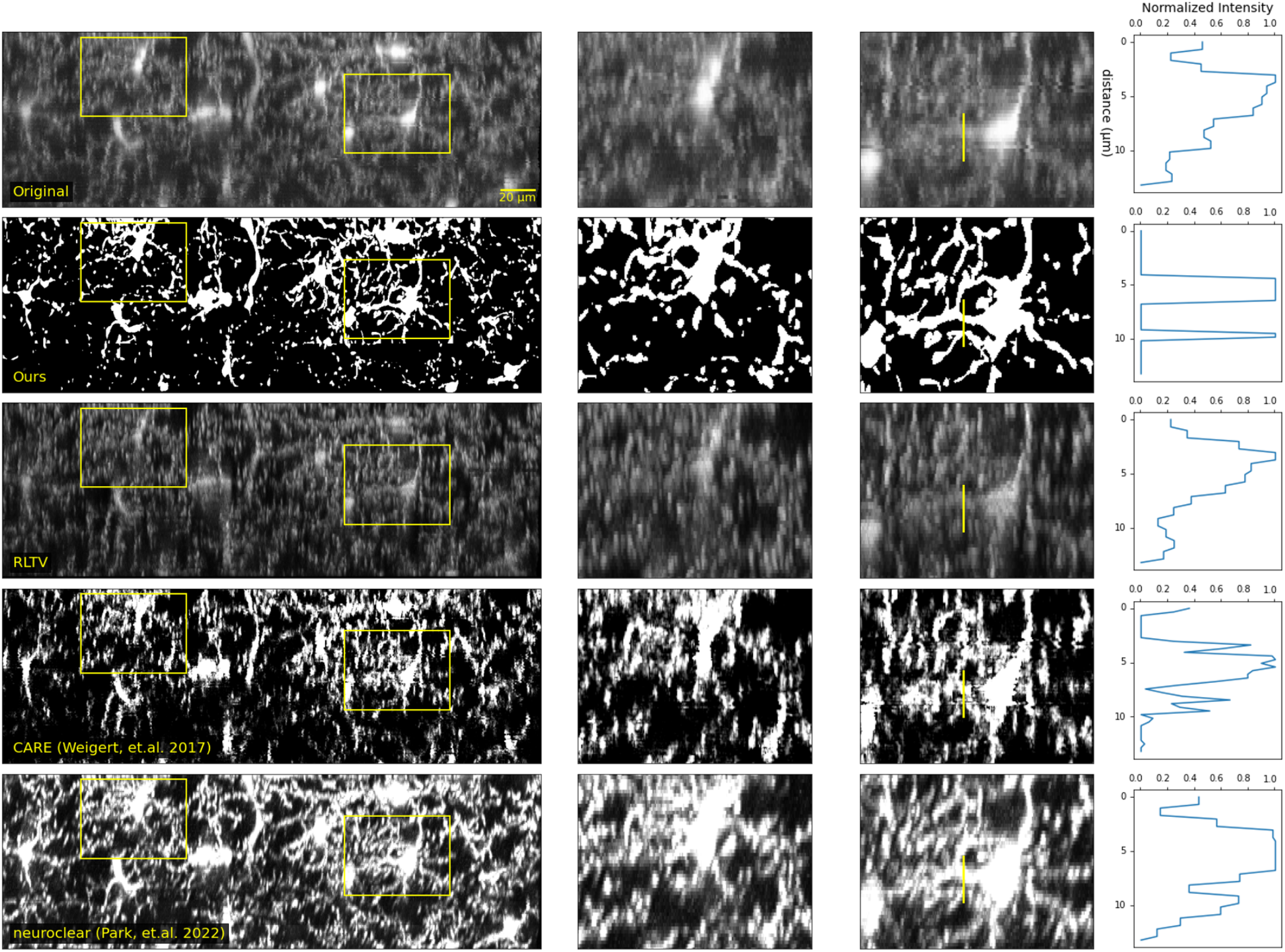
Results with actual two photon images of microglia. Far right: 2D MIP images. Second from right, second from left: Magnified views of the bounding boxes in the right image. Left: Intensity values along the yellow line in the second image from the left. Our model can clearly separate the two objects.

The model was able to reconstruct microglial processes. In the image on the right in Figure 3b, a structure that appeared to be a single process in the original image was detected as two processes after super-resolution and deblurring by the model.

## Discussion

We developed a method that simultaneously performs deblurring, super-resolution, and segmentation of two-photon microscopy images using unsupervised learning. We applied this model to simulation data and in vivo imaging of microglia and validated it.

As a result, the model demonstrated performance surpassing conventional methods. The results from the simulation data suggested that the model can accurately predict the shape of objects. When the model was applied to in vivo imaging of microglia, it successfully detected microglial processes that could not be detected by conventional methods.

Like other methods, our method is a model applied to images with fixed blur and voxel size. However, in deep imaging with two-photon microscopy, the blur increases proportionally with depth, and users may change the voxel size depending on the imaging target. As a future prospect, we plan to develop a model that can handle multiple resolutions and blur levels that increase with depth in a unified manner to accommodate such practical situations.

Conventional deblurring methods have suffered from issues such as the need for training data, low accuracy, and the inappropriate assumption of isotropy of the shape. Furthermore, segmentation and denoising had to be performed using separate methods, complicating the analysis process. The results in this paper suggest that our method solves the problems faced by conventional methods and enables the analysis of 3D shapes from two-photon microscopy images using unsupervised learning.

The application of our method is expected to provide insights into the dynamics of the 3D morphology of microglia in vivo, which has been difficult to analyze conventionally.

## Materials and Methods

### Simulation Setup

For the simulation images used in pre-training, we prepared 20 volumes of 1200 x 500 x 500 voxels and by randomly placing 1500 to 2500 objects (randomly selected from spheres, regular octahedrons, cubes, and lines) and deforming them with elastic deformation. These simulation images represent objects such as cells randomly distributed in tissue. Each object was assigned a brightness sampled from a log-normal distribution ℒ 𝒩 (*μ*_*B*_, *σ*_*B*_). In the fine-tuning experiments using simulations, we used images generated by randomly placing 200 to 300 objects in an image of the same size. For the blur model, we used a neural implicit function fitted to the PSF created using the parameters of the blur function intended for use in fine-tuning.

In the simulation images for pre-training, we shifted the position of the z-stacks based on the following equation of motion to reproduce the misalignment of images between zstacks:

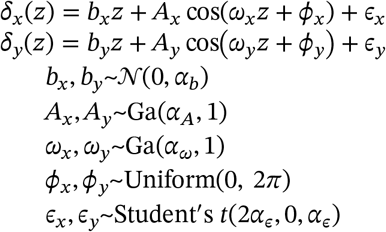

Where *δ* represents the position of the image, and *z* represents the index of the stack. The misalignment of the z-stacks was added within the data loader, and the parameters were adjusted so that the misalignment gradually increased during learning. Poisson noise was set as the noise for the simulation images.

### Mouse Brain Tissue Sample Preparation and Image Acquisition

Microglia were visualized in vivo in Cx3cr1^GFP/+^ mice (Jackson Laboratory). All images in this experiment were captured using a Nikon A1Plus microscope with a 20x objective lens, 2x digital zoom, a 0.17 mm correction lens, and a z-step of 1 μm.

### Image Preprocessing Steps

The images were normalized with a minimum value of 0 and a maximum value of 1.

### Validation Approach

IoU was used for validating the simulation data.

### Overview of Proposed Architecture

The proposed model consists of a combination of a deblurring model and a blur generation model. The deblurring model was trained in two stages. First, the model is pre-trained using simulation images created by the blur generation model. Next, the actual images are input to the model, and the two models are trained to minimize the error between the reconstructed blurred image generated by inputting the inference result into the blur generation model and the actual images (Figure 1a).

### Loss Formulation

During pre-training, the segmentation inference result 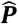 and brightness inference result 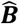 were compared with the label data ***P*** and ***B***. BCE loss 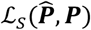 was used as the loss function for the segmentation inference result. For the brightness estimation result, the negative log-likelihood and the error from the prior distribution of brightness were used. Hereafter, the subscripts *i, j, k* represent the x, y, z coordinates of the voxel, respectively.

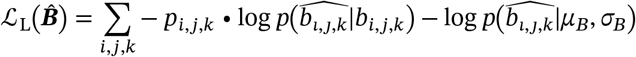

Overall loss function is as follows.

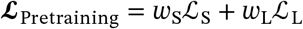

During fine-tuning, the deblurring model and the blur generation model are trained to minimize the reconstruction error. Denoting the deblurring model as *f*(***X***) and the reconstructed image by the combination of the deblurring model and the blur generation model as 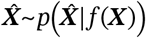, the reconstruction error ℒ_R_ is the negative log-likelihood loss of the Gaussian distribution between the original image ***X*** and the reconstructed image 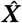,

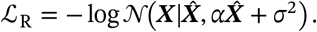

Here, α and σ are learnable.

In the finetuning process, the optimal strategy that the model can take is for each model to learn the identity transformation of the input image. In other words, it is easiest for models to learn to output the original image as it is for the deblurring model 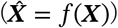 and to output the image as it is without blurring it as much as possible for the blur generation model 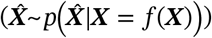 to minimize the reconstruction error. To prevent this, the following four losses were added.

#### VQ Loss

During fine-tuning, a vector quantization layer was set in the final layer of the deblurring model, and the output was binarized. Furthermore, the mean squared error ℒ_Q_ between the values before binarization ***P***^*cont*^ and the values after binarization ***P***^*bin*^ was added.

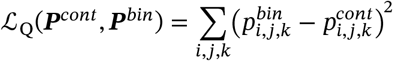

#### MRF Loss

To express the assumption that objects are clustered together, the following MRF loss ℒ_M_ was added to the binarized deblurring inference result.

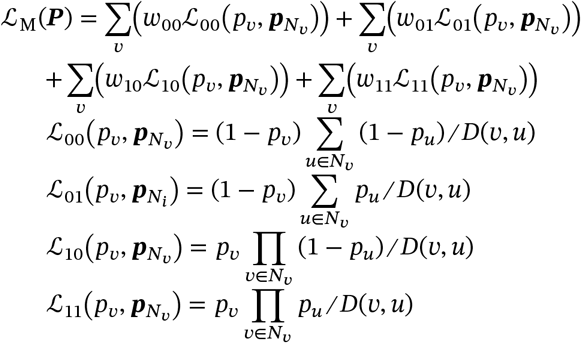

Here, *N*_*v*_ is the set of 26 neighbors of voxel *v. D*(*v, u*) represents the Euclidean distance between voxels.

#### EWC Loss

To avoid forgetting the learning results from the simulation data, EWC loss^xxxi^ was added to the deblurring model.

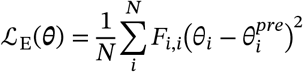

Here, ***θ***^*pre*^ is the mean value of the parameters of the deblurring model after pre-training, ***θ*** is the parameters of the deblurring model during reconstruction learning, and each has a total of N parameters.***F*** is the Fisher information matrix consisting of the following elements.

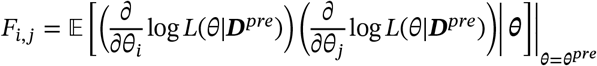

Where, ***D***^*pre*^ is the dataset used for pre-training.

#### PSF Loss

To reflect the specified PSF values, the mean squared error ℒ_P_ from the pre-set PSF 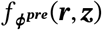 was added to the output value *f*_***ϕ***_(***r, z***) of the neural implicit PSF.

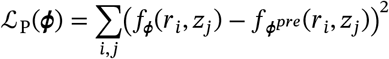

#### Luminance Loss

In addition to these loss functions, the ℒ_Lf_ (“Lum. Loss” in Figure 1a) is added to the brightness estimation result.

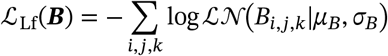

The final loss function is

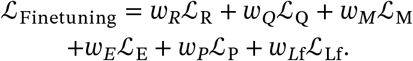

### Network Architecture

The deblurring model was created based on a 3D U-Net^xxxii^ structure. The model consists of an initial layer, U-Net layers, and a final layer.

The initial layer consists of a brightness adjustment hill function, upsampling by trilinear interpolation, and a 7×7×7 convolutional layer.

The U-Net layers are composed of a Contraction path and an Expanding path, each consisting of 5 stacked blocks. Each layer of the Contraction path consists of Residual blocks and downsampling (MaxPooling and 1×1×1 convolution, activation function). Each layer of the Expanding path consists of Residual blocks and upsampling (trilinear interpolation and 1×1×1 convolution, activation function). For each Residual block, with reference to the Fully Pre-activation ResNet block^xxxiii^, a structure of batch normalization - activation function - convolutional layer - batch normalization - activation function - dropout - convolutional layer was adopted.

The final layer is divided into a shape estimation layer and a brightness estimation layer. Both layers are input from the final output of the UNet layer. The common structure consists of Residual blocks, a final convolutional layer (3x3x3) that outputs 1 channel, and an activation function (sigmoid function). The number of blocks in the Residual block is set to 2. During reconstruction learning with actual images, a quantization layer follows the shape estimation layer, and a process is performed to set pixels with brightness values above a threshold to 1 and others to 0. To make this process differentiable, the stop gradient estimator proposed in VQVAE was used^xxxiv^.

The blur generation model consists of 1) a model of the process of fluorescence emission from objects, 2) a model of the process of fluorescence blurring, and 3) a process of noise being added during imaging.

1. The process of fluorescence emission from objects is assumed to be independent for each voxel and is expressed in the following form of the Hadamard product.

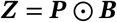

***Z*** can be regarded as the fluorescence intensity in the absence of blur.

2) The process of fluorescence being blurred and captured is assumed to be independent for each voxel, and the PSF ***f*** is convolved with the fluorescence intensity of each voxel in the absence of blur.

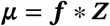

A neural implicit function was used as the PSF. The neural implicit function takes coordinates consisting of the horizontal distance and vertical distance from the center of the PSF as input and outputs the corresponding PSF value. The network architecture consists of a batch normalization layer, a linear layer that projects the 2-dimensional input coordinates, a sigmoid activation function, another batch normalization layer, a linear layer that projects to a single output, and a final sigmoid activation function. The network is trained with input coordinate data and corresponding correct PSF values. In all experiments in this paper, the Gibson-Lanni model was used as the correct PSF.

3) For image noise, Gaussian distribution noise with variance *α****μ*** + *σ*2 was assumed.

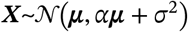

### Data Preparation

In all training, the model was provided with randomly cropped regions of (240, 112, 112).

The microglia data used for learning and inference contain regions without objects. With such sparse data, the optimal model output may always be zero regardless of the input. To prevent this, the model was supplied with data using the following procedure to provide small regions containing as many objects as possible.

A random small region is obtained, its mean brightness value is taken, and when inputting data to the model, *r* is sampled from a uniform distribution, and the data is input to the model only if the mean value exceeds *r*; otherwise, the small region and *r* are resampled.

For data augmentation, 90-degree rotation, left-right flipping, and masking (randomly selecting a (1, 10, 10) region of the input and setting it to zero) were used.

### Deblurring Network Architecture Details

The number of channels in the UNet layers was set to [16, 32, 64, 128, 256]. Downsampling and upsampling were performed by a factor of 2. The number of blocks in each layer of each block was set to 2. ReLU was used as the activation function. The dropout rate was set to 0.5.

### Blur Generation Model Architecture Details

In all experiments, the Gibson-Lanni model was used as the label data for the PSF. The number of channels in the middle layer of the neural implicit function PSF was set to 40, and sigmoid was used as the activation function. FFT convolution (Fast Fourier-Transformation) was used for the convolution of the PSF and the image.

### Hyperparameters for Training

The batch size was set to 1, and the number of small regions loaded per epoch was set to 200. In all experiments, *w*_*S*_= 1, *w*_*L*_ = 1 were set during pre-training. For fine-tuning on the simulation images in Figure 2, *w*_*R*_ = 1, *w*_*Q*_ = 1, *w*_*M*_ = 1, *w*_00_= −0.5, *w*_01_ = 0.5, *w*_10_ = 1, *w*_11_ = −1, *w*_*Lf*_ = 1, *w*_*E*_ = 0.1, *w*_*P*_ = 10 were set. For fine-tuning in Figure 3, only *w*_*M*_ = 2 was changed.

The misalignment parameters *α*_*b*_, *α*_*A*_, *α*_*ω*_ were linearly increased from 0 to 0.2, 2, 2, respectively. *α*_*λ*_ was linearly decreased from 10 to 2. This process was performed over 60 steps.

The noise parameters were set to *α* = 0.1,and *σ* = 0.01 The Gibson-Lanni model parameters were set to NA = 0.3, λ_wavelength_ = 0.5 nm, M = 25, RIspecimen = 1.4, RIcoverslip = 1.5, RIimmersion = 1.33, ti = 150 μm, tg = 170 μm, pZ = 0 in all experiments.

In all experiments, the deblurring model before pre-training and the neural implicit PSF were initialized using Gaussian initialization.

### Optimization

Deblurring model was trained using the Adam optimizer^xxxv^ with a learning rate of 10^−3^ during pre-training and 10^−3^ during fine-tuning. Warmup was linearly increased from 0 to the learning rate over 10 epochs. Reduce on plateau was used with decay 0.1 and patience 20 epochs. In all training, the total number of epochs was 200, and the weights with the smallest validation loss were used.

The neural implicit PSF was trained by full-batch learning using the RProp optimizer^xxxvi^ with a learning rate of 10^−3^ and optimized until the error reached 10^−5^

### Machine Used for Experiments

The experiments were performed on a computer equipped with an NVIDIA GeForce RTX 3090 graphics card and an AMD Ryzen 9 5900X 12-Core Processor CPU.

